# Making memories last: The peripheral effect of direct current stimulation on strengthening memories

**DOI:** 10.1101/2022.07.06.498966

**Authors:** Alison M. Luckey, S. Lauren McLeod, Yuefeng Huang, Anusha Mohan, Sven Vanneste

## Abstract

Most memories that are formed are forgotten, while others are retained longer and are subject to memory stabilization. We show that non-invasive transcutaneous electrical stimulation of the greater occipital nerve (NITESGON) using direct current during learning elicited a long-term memory effect. However, it did not trigger an immediate effect on learning. A neurobiological model of long-term memory proposes a mechanism by which memories that are initially unstable can be strengthened through subsequent novel experiences. In a series of studies, we demonstrate NITESGON’s capability to boost the retention of memories when applied shortly before, during or shortly after the time of learning by enhancing memory consolidation via activation and communication in and between the locus coeruleus pathway and hippocampus by modulating dopaminergic input. These findings may have a significant impact for neurocognitive disorders that inhibit memory consolidation such as Alzheimer’s disease.

## Introduction

Research on enhancing and preserving human memory has substantially increased in the last few decades, due in large part to the prevalence and inexorable condition of Alzheimer’s disease. Recent investigations have begun to assess the perspective clinical significance of therapeutic non-invasive brain stimulation techniques to modify neuroplasticity and upregulate neuronal excitability in different neurological conditions including memory deficits^1^. There is presently an ongoing debate whether non-invasive electrical stimulation of the scalp modulates the excitability of neurons directly^2, 3^. Interestingly, a series of experiments in rats and humans isolated the transcranial and transcutaneous mechanisms of non-invasive electrical stimulation and showed that the reported effects are mainly caused by transcutaneous stimulation of peripheral nerves^4^. Similarly, it was demonstrated that nerve stimulation paired with an auditory or motor task can induce targeted plasticity in animals^5, 6^. Our recent work demonstrated that non-invasive transcutaneous electrical stimulation of the greater occipital nerve (NITESGON) using direct current during learning induces improvements in memory recall in younger (18–25 years) and older (>65 years) adults up to 28 days after learning^7, 8^. Intriguingly, NITESGON yields a long-term memory effect, but did not trigger an immediate effect on learning, suggesting that the effect is generated during the consolidation of memories^8, 9^, as opposed to the during learning or encoding of new memories.

Most episodic-like memories that are formed are forgotten, while others are retained for longer periods of time and are subject to memory stabilization^10, 11, 12^. This is referred to as synaptic consolidation, a process which stabilizes new information into memory over a timespan of minutes to hours. The neurobiological account of synaptic consolidation has proposed a synaptic tag-and-capture mechanism whereby new memories that are initially weak and unstable are tagged to be captured by late-phase long-term potentiation (LTP) to become stable^13, 14^. This mechanism can explain how weak behavioral training that would typically be forgotten will consolidate when followed by a novel behavioral experience − an effect referred to as behavioral tagging^15^. The neural mechanism that controls this novelty response is the LC-NA pathway^16, 17^. Animal research further indicates that direct electrical stimulation of the LC modulates hippocampal synaptic consolidation^18–20^. We hypothesize that NITESGON modulates projections to the hippocampus via the LC-NA system and induces memory stabilization by modulating synaptic consolidation in the hippocampus via the mechanism of behavioral tagging^8^.

The present study tests the hypothesis if NITESGON induces a long-term memory effect by strengthening memories via behavioral tagging across the span of eight experiments. The first set of experiments aims to confirm the behavioral tagging hypothesis as the potential mechanism inducing memory consolidation via NITESGON, whereby the second set of experiments examines the underlying brain network involved in synaptic consolidation and investigates the underlying neural mechanism that is associated with behavioral tagging induced by NITESGON.

### Experiment 1. NITESGON during or immediately after training

The idea behind behavioral tagging suggests that weak memories that are regularly unstable and likely to be forgotten will solidify following a novel experience^15^. That is, consolidation is facilitated by applying a strong stimulus alongside a weak stimulus within a critical time window. Recent research revealed a direct link between the LC and behavioral tagging, attributable to the pivotal role the LC plays during the presentation of a salient or arousing event (i.e., strong stimulus)^21, 22^, as well as being at the helm of regulating the synthesis of new proteins required for memory consolidation in the hippocampus^23^. Furthermore, studies have shown modulation of memory consolidation with increases in stress and arousal that are mediated via the LC pathway^19, 20^. Moreover, animal research has indicated that direct electrical stimulation of the LC modulates hippocampal synaptic transmission fundamental for memory consolidation^18^.

Seeing that NITESGON activates the LC pathway, which plays an important role in memory consolidation, we hypothesize that participants will be able to establish long-term memories upon modulating the LC both during learning, as shown before, as well as immediately after learning. This would directly test if NITESGON plays a more central role during encoding or the consolidation phase. To test this hypothesis, participants learned a word association task and were tested 7 days later, on how many word-associations they were able to correctly recall. Active or sham NITESGON was applied via electrodes placed over the left and right C2 nerve dermatome at a constant current of 1.5 mA either during or immediately after learning the word-association task on visit 1.

To further explore the effect of NITESGON, resting-state EEG (rsEEG) and salivary α-amylase (sAA) were collected immediately before and after NITESGON on visit 1. Previous research has revealed an increase in sAA, a marker of endogenous NA activity, immediately following NITESGON^8, 24^. Furthermore, previous investigations have demonstrated that LC discharge enhances synchronization of gamma activity in the hippocampus in rats^25^ and have exhibited gamma oscillations’ critical role in long-term memory formation and potential to predict subsequent recall^26, 27^. Based on these findings, we hypothesized that NITESGON would induce an increase in sAA as well as gamma activity in the medial temporal lobe that will correlate with successful recall during the second visit 7 days after learning the task.

On visit 1, no difference was observed regarding the number of word-associations learned between the three condition groups (i.e., sham NITESGON during learning and after learning, active NITESGON during learning and sham NITESGON after learning, or sham NITESGON during learning and active NITESGON after learning) (*F* = .24, *p* = .79; see fig. 1a), thus indicating that NITESGON had no effect on learning the word-association task. Results revealed a significant difference in memory recall 7 days after NITESGON was applied either during learning (46.09 ±15.06%, *p* = .012) or after learning the task (47.65 ±13.27%, *p* = .005) relative to the sham condition employed during both learning and immediately after learning the task (33.38 ±12.57%) (*F* = 5.24, *p* = .009; see fig 1b). However, no difference was attained on recall 7 days later between the conditions of NITESGON applied during learning the task or immediately after learning the task (*p* = .75). A significant increase in sAA (*F* = 7.69, *p* = .010; see fig. 1c) was revealed during learning (before: 88.79 ±50.48 vs. after: 149.82.6 ±82.67; *p* < .001) and after learning in comparison to the sham group (before: 100.28 ±41.95 vs. after: 114.30 ±41.02; *p* = .14). Memory recall 7 days later correlated with the difference in sAA levels on visit 1 (pre vs. post) (r = .59, p < .001; fig. 1d). Memory recollection 7 days after stimulation was associated with increased gamma power in the medial temporal cortex as well as the precuneus and dorsal lateral prefrontal cortex immediately after stimulation (average R^2^ = .11, p = .011; see fig. 1e).

**Figure 1.**
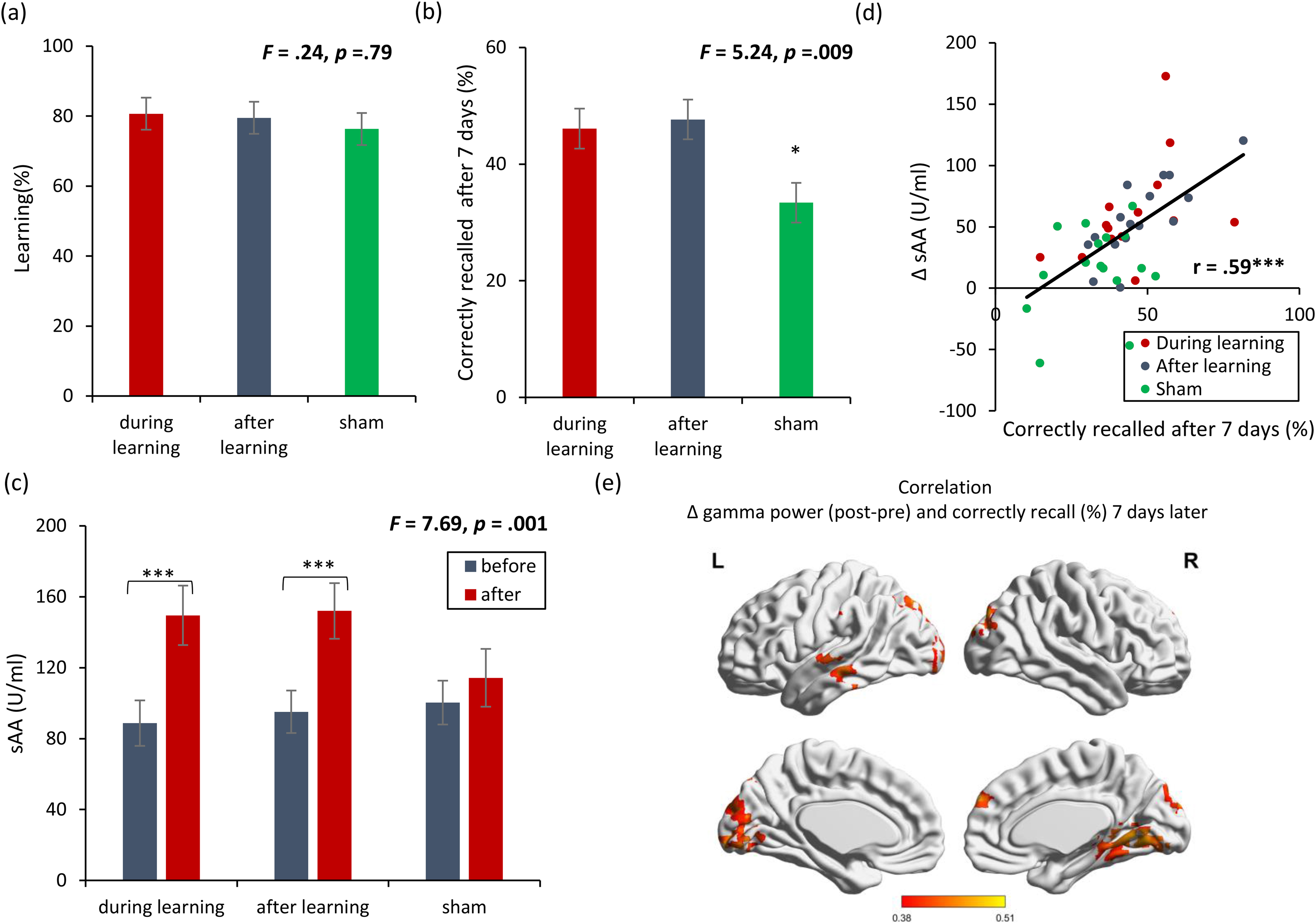
NITESGON immediately after training can enhance memory. (a) No difference was observed in the cumulative learning rate between active and sham NITESGON during or immediately after the study phase of the word association memory task. (b) NITESGON during or immediately after the word association memory task can improve memory recall 7-days after the study phase for the active relative to the sham group. (c) After NITESGON sAA levels increase for both active groups, but not for sham NITESGON. (d) Memory recall 7-days later correlates with the difference in sAA levels during the first visit (pre vs post study phase). (e) Improved memory recall 7-days after stimulation is associated with increased activity in the medial temporal lobe as well as anterior and posterior cingulate cortex immediately after NITESGON for the gamma frequency band. Error bars, s.e.m. Asterisks represent significant differences (* *p* < .05; ** *p* < .01).

### Experiment 2. NITESGON during second task – retroactive strengthening of memories

Experiment 1 suggests that NITESGON generates an effect during the consolidation phase as opposed to the learning-encoding phase due to no effect of NITESGON being exhibited during learning between the different groups, but both stimulating during or after learning the task induced a long-term memory effect. Bearing in mind the definition of behavioral tagging that indicates that the pairing of a strong stimulus and a weak stimulus within a critical time window can induce memory stabilization of the weak stimulus, NITESGON can be seen as the mechanism that induces a similar action as a strong stimulus, and through the mechanism of behavioral tagging strengthen the weak stimulus (i.e., the word-association task).

Prior research on behavioral tagging has shown that items paired with an electric shock (i.e., Pavlovian fear conditioning task) had a retroactive memory effect on items learned before the fear conditioning task. This provided evidence for a generalized retroactive memory enhancement, whereby information can be retroactively credited as relevant, and therefore remembered^15^. Interestingly, LC activation occurs in close relation to the intensity of the Pavlovian behavior^28^. Hence, to explore the effect of the LC on behavioral tagging, we verified if NITESGON applied during a second task would result in a significant retroactive memory effect on the first task as predicted by behavioral tagging.

To test the hypothesis, Experiment 2 had participants take part in a word-association task followed by a spatial navigation object-location task while receiving active or sham NITESGON during the second task. These two types of tasks were selected because they would not interfere with one another seeing that both require different episodic information. rsEEG data and sAA were collected immediately before and after the two tasks on visit 1. Two memory tests were taken 7 days after learning the word-association and spatial navigation tasks.

On visit 1, no difference in learning (*F*= .32, *p* = .73; see fig. 2a) was observed for both the first (*F*= .09, *p* = .98) and second (*F* = .64, *p* = .43) tasks between the active and sham NITESGON groups. On visit 2, 7 days after initial learning, a significant effect was obtained for recall (*F*= .6.82, *p* = .007; see fig. 2b) for both the first (*F* = 6.28 *p* = .022) and second tasks (*F* = 7.51, *p* = .013), revealing an increase in word recall (46.26 ±3.76% vs. 37.88 ±9.88%), as well as object-location recall (51.82 ±7.75% vs. 44.39 ±3.68%) for the active group in comparison to the sham group. Furthermore, a significant increase in sAA (*F* = 7.44, *p* = .014; see fig. 2c) was revealed in the active group (before: 74.01 ±26.58 vs. after: 107.01 ±25.98; *p* < .001) in comparison to the sham group (before: 60.61 ±37.93 vs. after: 69.61 ±35.07; *p* = .18). This increase in sAA correlated with how many items they recalled 7 days after the learning phase for both the word-association task (r = .52 *p* = .019; see fig. 2d) and the object-location task (r = .57, *p* = .008; see fig. 2e). Memory recollection 7 days after stimulation was associated with increased gamma power in the medial temporal cortex immediately after stimulation for both the first (r = .41, *p* = .009; see fig. 2f) and second memory tasks (r = .35, *p* = .018; see fig. 2g).

**Figure 2.**
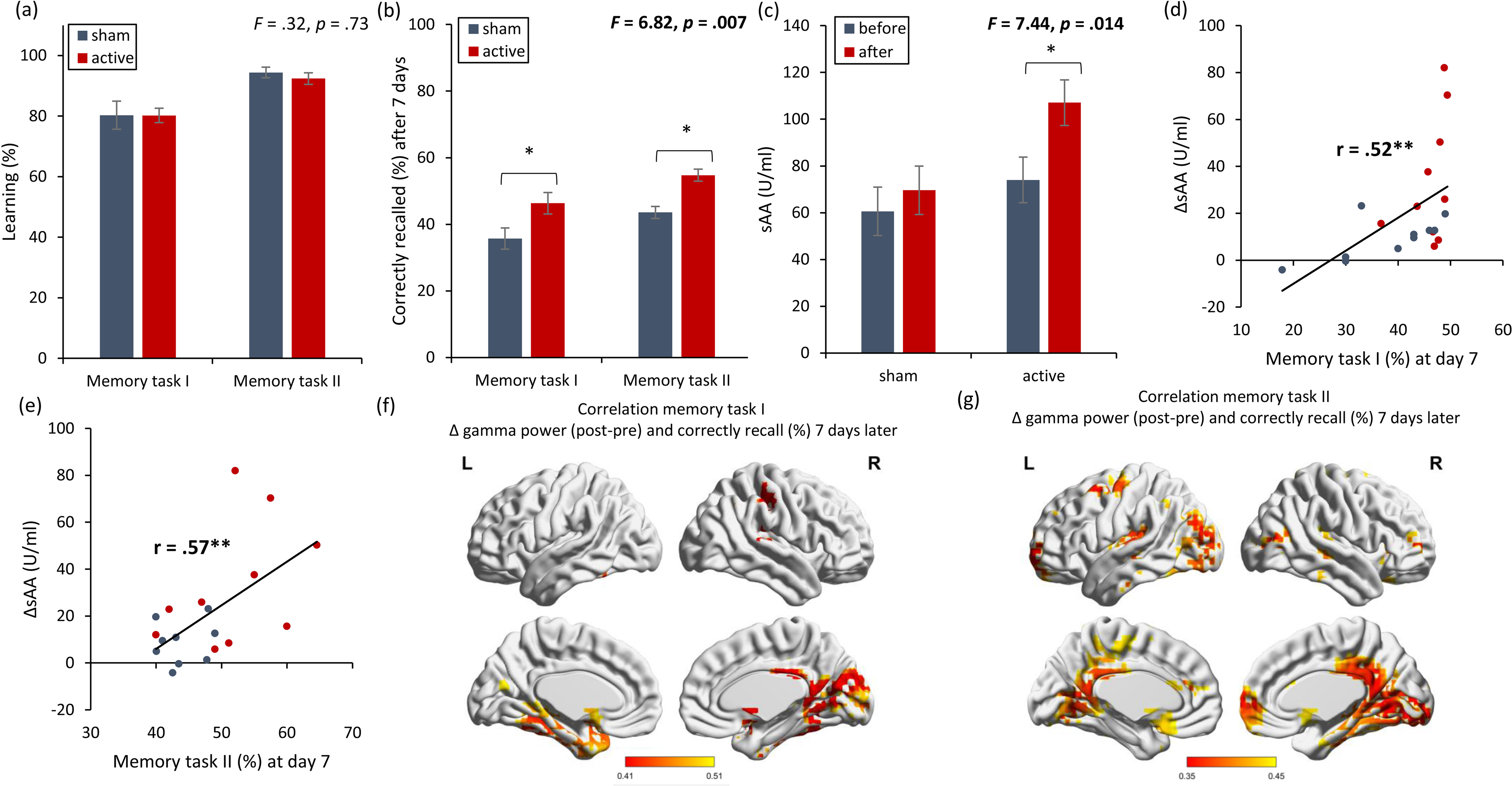
NITESGON has a retroactive memory effect – NITESGON during the second task. (a) No difference was observed in the cumulative learning rate between active and sham NITESGON after the study phase for the first task (i.e., word-association task) or second task (i.e., object-location task). (b) NITESGON can improve memory recall 7-days after the study phase for the active relative to the sham group for both the first and second tasks. (c) After NITESGON sAA levels increase for active group, but not for sham NITESGON. (d,e) Memory recall 7-days later correlates with the difference in sAA levels during the first visit (pre vs post study phase) for the first and second tasks. (f,g) Improved memory recall 7-days after stimulation is associated with increased activity in the medial temporal lobe immediately after NITESGON for the gamma frequency band. Error bars, s.e.m. Asterisks represent significant differences (* *p* < .05; ** *p* < .01).

### Experiment 3. NITESGON during first task - proactive strengthening of memories

Experiment 2 revealed a retroactive memory effect 7 days after initial learning for the active NITESGON group in comparison to the sham NITESGON group, fitting well with the behavioral tagging hypothesis. In addition to a retroactive memory effect, previous research on behavioral tagging also revealed items paired with an electric shock had a proactive memory effect, whereby items learned after the fear conditioning task were remembered^15^. Here, we conducted the exact same experiment as in Experiment 2 but applied NITESGON during the first task and not during the second task to test the hypothesis if NITESGON can induce a proactive memory effect on the second task although we stimulate during the first task. This would further support the hypothesis that NITESGON induces a long-term memory effect via the mechanism of behavioral tagging through activation of the LC pathway.

On visit 1, no significant difference (*F* = 2.26, *p* = .13; see fig. 3a) was found between the active and sham groups regarding how many words or objects participants learned for both the first task (i.e., word-association task) (*F* = 1.60, *p* = .22) and the second task (i.e., object-location task) (*F* = 3.30, *p* = .08). During the second visit, 7 days after learning the tasks, participants that received active NITESGON (*F* = 4.66, *p* = .021; see fig. f3b) recalled more words for the first task (i.e., word association task) (*F* = 6.32, *p* = .020) and the second task (i.e., object-location task) (*F* = 4.87, *p* = .038) than those who received sham NITESGON, indicating that the active NITESGON group (44.27 ±10.97) showed significant improvement in comparison to the sham NITESGON group (30.46 ±15.13) for the word-association task. For the object-location task, the active NITESGON group (53.33 ±11.29) demonstrated a significant increase in the number of correctly recalled objects-locations than the sham NITESGON group (44.17 ±7.75). Our data revealed that there was a significant interaction effect for sAA (*F* = 4.66, *p* = .021; see fig. 3c), denoted by the active group’s increase in sAA (123.10 ±43.63) in comparison to the sham group (91.38 ±44.67) (*F* = 4.53, *p* = .039) immediately after learning. No significant difference (*F* = .012, *p* = .91) was obtained in sAA for the active group (78.72 ±47.67) in comparison to the sham group (76.77 ±39.87) before learning the association tasks. This increase in sAA seen in the active group correlated with how many items they recalled 7 days after the learning phase for both the word-association task (r = .42, *p* = .039; see fig. 3d) and the object-location task (r = .51, *p* = .012; see fig. 3e). Memory recollection 7 days after stimulation was associated with increased gamma power in the medial temporal cortex immediately after stimulation for both the first (r = .32, *p* = .037; see fig. 3f) and second memory tasks (r = .52, *p* = .012; see fig. 3g).

**Figure 3.**
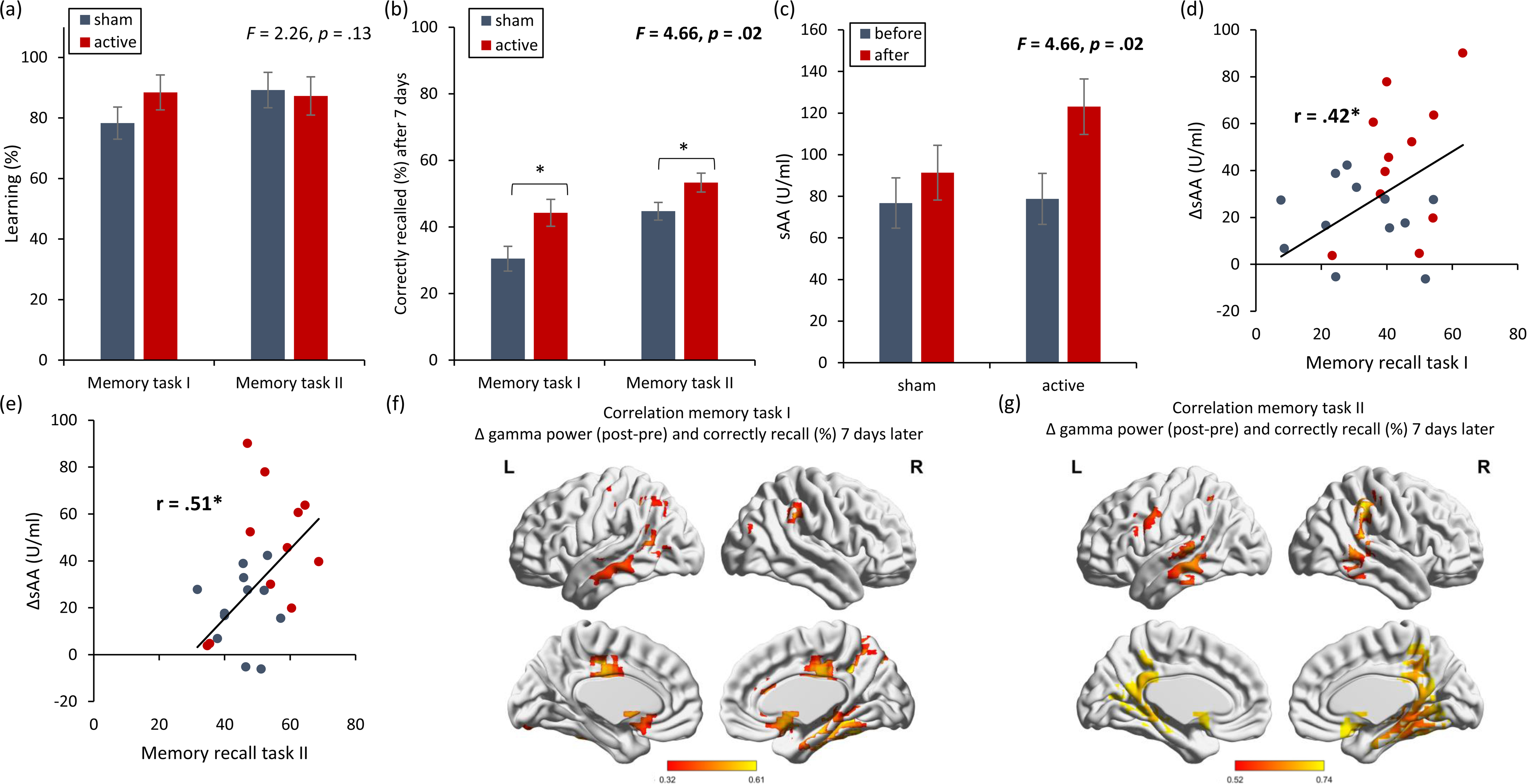
NITESGON has a proactive memory effect – NITESGON during the first task. (a) No difference was observed in the cumulative learning rate between active and sham NITESGON after the study phase for the first task (i.e., word-association task) or second task (i.e., object-location task). (b) NITESGON can improve memory recall 7-days after the study phase for the active relative to the sham group for both the first and second tasks. (c) After NITESGON sAA levels increase for active group, but not for sham NITESGON. (d,e) Memory recall 7-days later correlates with the difference in sAA levels during the first visit (pre vs post study phase) for the first and second tasks. (f,g) Improved memory recall 7-days after stimulation is associated with increased activity in the medial temporal lobe immediately after NITESGON for the gamma frequency band. Error bars, s.e.m. Asterisks represent significant differences (* *p* < .05; ** *p* < .01).

### Experiment 4. NITESGON during interference task

Experiment 2 and 3 revealed both retroactive and proactive memory effects 7 days after initial learning of the two tasks. To further explore if NITESGON is linked to behavioral tagging, we introduced a learning task similar to the Swahili-English verbal associative task in Experiments 1, 2 and 3. Considering how new memories are susceptible to interference immediately after its encoding^29, 30^, it is believed that conducting two consecutive, like-minded word-association (i.e. Swahili-English and Japanese-English) tasks will result in one’s consolidation process interfering with that of the other^31^. Considering how our previous experiments suggests the effect obtained by NITESGON improves the consolidation of information via behavioral tagging, it is possible that NITESGON on the first task might help reduce the overall interference effect on the second task.

To test the hypothesis, participants participated in two separate word association tasks (i.e.., the Swahili-English and Japanese-English; the order of tasks was randomized across participants) while receiving either active or sham NITESGON during the first task. rsEEG data and sAA were collected immediately before and immediately after the two tasks on visit 1. Two memory tests were taken 7 days after learning both word-association tasks.

On visit 1, no significant difference (*F* = .84, *p* = .37; see fig. 4a) was detected for learning during the first (*F* = .27, *p* = .61) and second task (*F* = .01, *p* = .94) between the active and sham NITESGON groups. Seven days later, during the recall phase, we found a significant interaction effect for recall (*F* = 4.97, *p* = .034; see fig. 4b). For both the first (*F* = 3.67, *p* = .048) and the second tasks (*F* = 7.89, *p* = .009), a significant increase in number of words correctly recalled was observed in the active (first task: 35.51 ±8.68; second task: 34.76 ±11.74) compared to the sham group (first task: 29.80 ±9.72; second task: 24.64 ±8.10). The active group displayed no significant difference between the first and the second task in how many words partipicant were able to recall (difference: .76 ±4.93) (*F* = .29, *p* = .60), while the sham group demonstrated an interference effect of the first task on the second task (difference: 5.16 ±5.99) (*F* = 14.11, *p* = .001).

**Figure 4.**
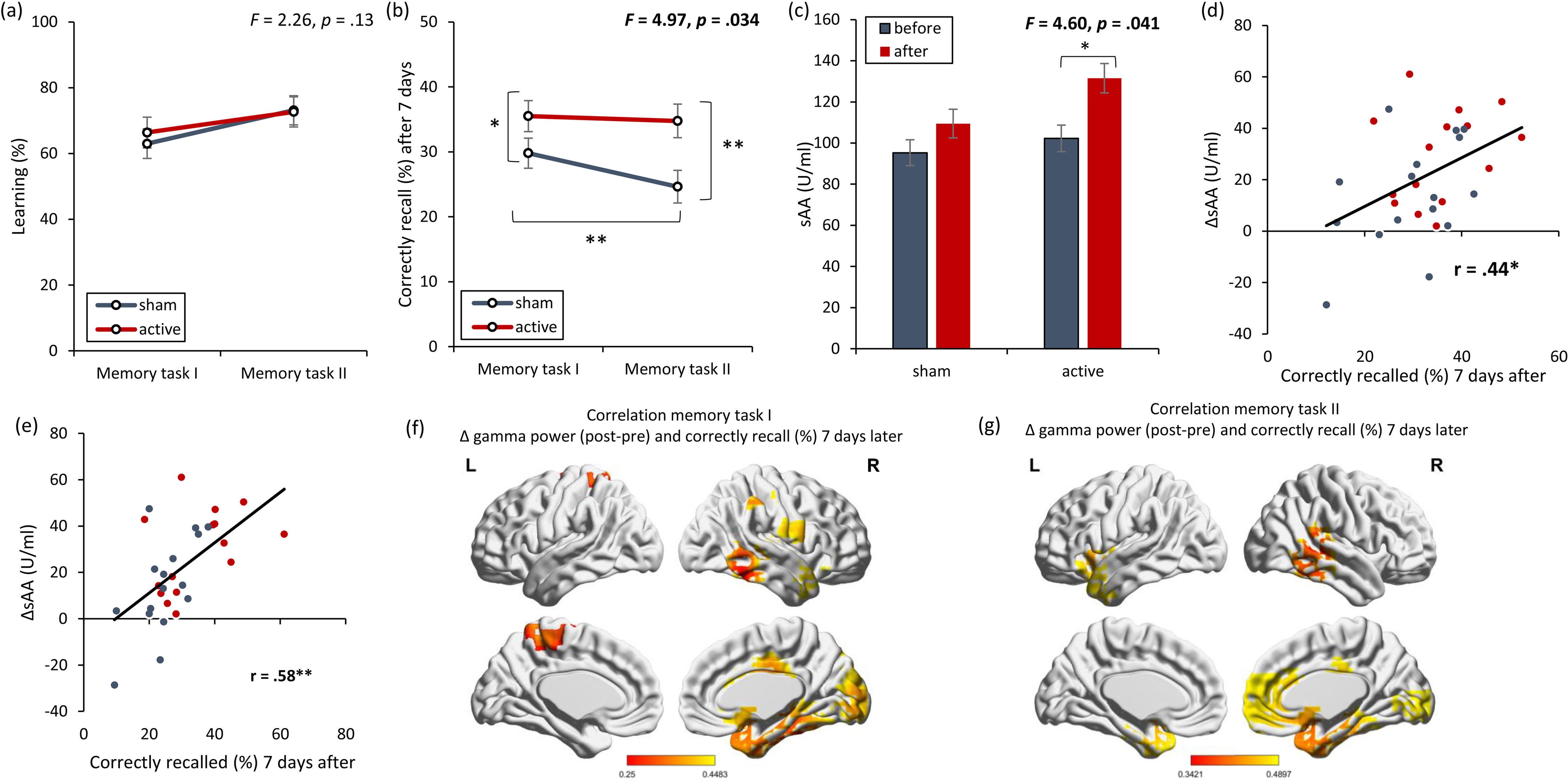
NITESGON reduces the interference effect. (a) No difference was observed in the cumulative learning rate between active and sham NITESGON after the study phase for the first task (i.e., word-association task) or second task (i.e., word-association task). (b) NITESGON can improve memory recall 7-days after the study phase revealing that the interference effect is reduce for the active relative to the sham group for both the first and second tasks. (c) After NITESGON sAA levels increase for the active group, but not for sham NITESGON. (d,e) Memory recall 7-days later correlates with the difference in sAA levels during the first visit (pre vs post study phase) for the first and second tasks. (f,g) Improved memory recall 7-days after stimulation is associated with increased activity in the medial temporal lobe immediately after NITESGON for the gamma frequency band. Error bars, s.e.m. Asterisks represent significant differences (* *p* < .05; ** *p* < .01).

Our data revealed that there was a significant interaction effect for sAA (*F* = 4.60, *p* = .041; see fig. 3c). An increase in sAA was observed for the active group (123.10 ±43.63) in comparison to the sham group (91.38 ±44.67) (*F* = 4.53, *p* = .039) immediately after the learning. However, no significant difference (*F* = .012, *p* = .91) was obtained in sAA for the active group (78.72 ±47.67) in comparison to the sham group (76.77 ±39.87) before learning the word-association tasks. This increase in sAA correlates with how many words they recalled 7 days after the learning phase for both the first word-association task (r = .44 *p* = .014; see fig. 4d) and the second word-association task (r = .58, *p* = .001; see fig. 4e).

Memory recollection 7 days after stimulation was also associated with increased gamma power in the medial temporal cortex immediately after stimulation for both the first (r = .25, *p* = .042; see fig. 4f) and second memory tasks (r = .34, *p* = .032; see fig. 4g).

### Experiment 5. Effect of NITESGON is not sleep dependent

Our behavioral experiments suggest that NITESGON targeting the LC is involved in synaptic consolidation via the behavioral tagging mechanism. It is assumed that synaptic consolidation occurs over a timespan of minutes to hours after encoding the information, thus this effect is time-dependent^32^. Furthermore, prior research has revealed that retroactive memory enhancement (i.e., evidence for behavioral tagging) emerges within 6 hours and is not dependent on sleep^15^. Based on these previous findings and the assumption that NITESGON modulates synaptic consolidation via the mechanism of behavioral tagging, we hypothesize that sleep would not mediate the memory effect induced by NITESGON.

Experiment 5 compared two groups of participants undertaking a word-association task paired with active NITESGON. One group of participants slept between the word-association task at 8 p.m. and the test phase the next day at 8 a.m., whereas the other group did not sleep between the learning phase at 8 a.m. and the test phase that took place at 8 p.m. that same day. A comparison between the two groups revealed no significant difference in the number of words learned during the learning phase (*F* = .26, *p* = .62; see fig. 5a) as well as no significant difference between the two groups when tested 12 hours later (*F* = .31, *p* = .59; see fig. 5b). Participants who slept in-between the learning and test phase correctly recalled 89.99 ±13.09% of word pairs and participants who did not sleep in-between the learning and test phase correctly recalled 87.23 ±7.41% of word pairs.

**Figure 5.**
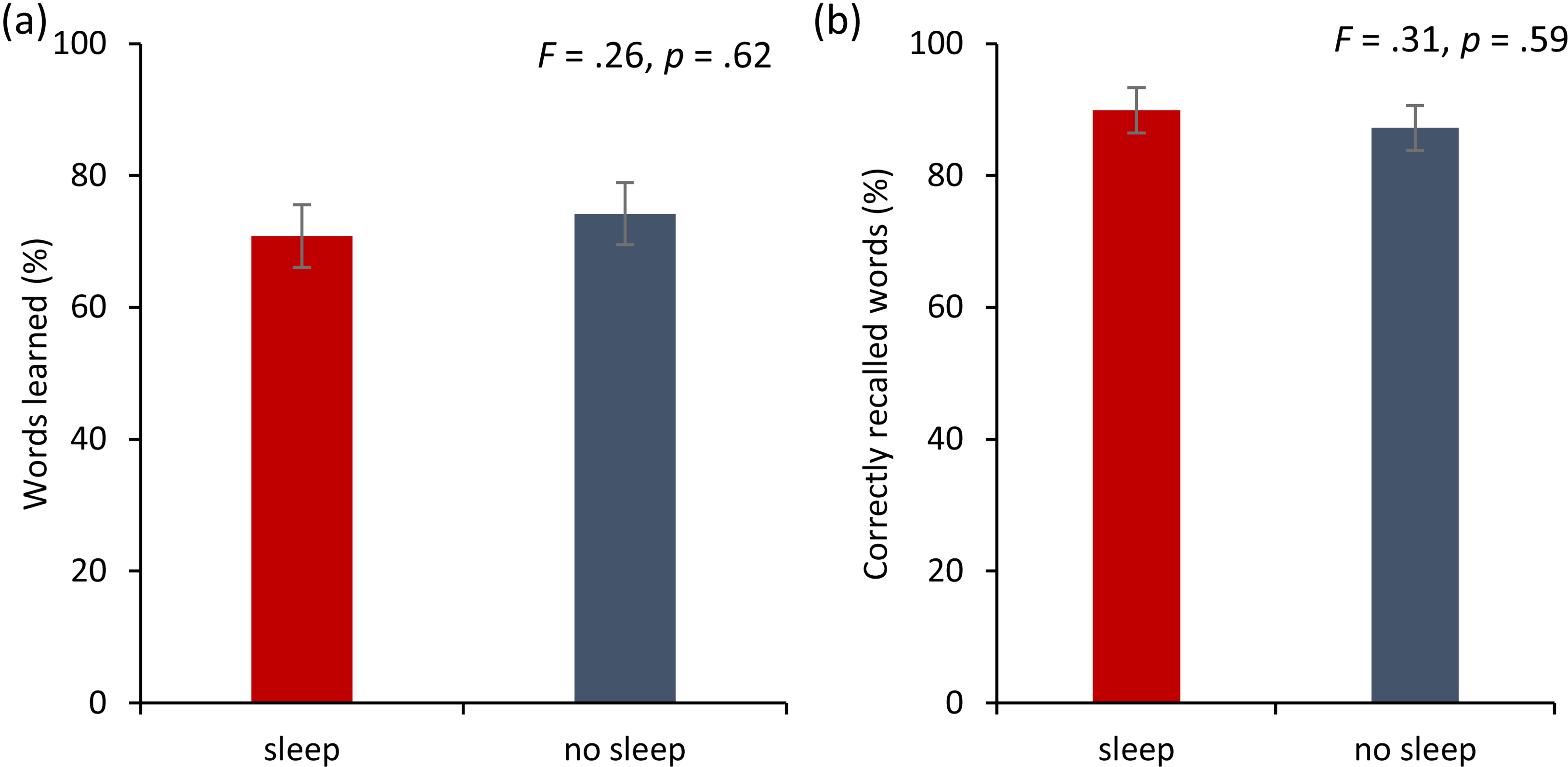
NITESGON and sleep. (a) No difference was observed in the cumulative learning rate between participants who had slept versus those who had not slept after NITESGON applied during the study phase. (b) Sleep has no effect on memory recall 12-hours after the study phase. Error bars, s.e.m. Asterisks represent significant differences (* *p* < .05; ** *p* < .01).

### Experiment 6. LC – Hippocampus activity & connectivity

In addition to our behavioral experiments confirming the hypothesis that NITESGON targeting the LC is involved in memory consolidation via the behavioral tagging mechanism, Experiment 5 revealed that sleep does not mediate the effect generated by NITESGON. Here, in the second set of experiments, we explored the brain network modulated by NITESGON and investigated the potential underlying neural mechanism.

The hippocampus is the key brain area that has been associated with synaptic memory consolidation. This area receives neuromodulatory input from multiple brain regions that regulate synaptic plasticity, such as, the LC and the ventral tegmental area (VTA). Both brain areas enhance retention of everyday memories in the hippocampus^17^. Moreover, animal research identified the VTA and the LC as regulators of hippocampal dependent long-term memory formation due to their role in regulating the synthesis of new proteins required during the behavioral tagging process, therefore allowing for the consolidation of lasting memories^23^. However, recent studies have shown that the VTA projections to the hippocampus are scarce, while the LC projections are abundant^17, 33, 34^. Therefore, the VTA may only play a limited role in late-phase LTP^17, 33, 34^, whereas the LC is conceivably the primary source of synaptic modulation responsible for tuning cells in the hippocampus^33^. Additionally, several previous observations have shown electrical and pharmacological stimulation of the LC modulated hippocampal synaptic transmission^18, 35^, whilst modulation of the VTA did not significantly mediate synaptic transmission but rather suggest a greater role in salience and motivational drive underlying emotion-based learning^33, 35^.

We conducted a resting-state functional connectivity MRI study to verify the relationship between changes in the LC and hippocampus as well as the VTA and hippocampus. We hypothesized that participants who received active NITESGON would show increased activity in the LC and hippocampus, but not in the VTA, and increased functional connectivity between the LC and hippocampus, but not between the VTA and hippocampus. We scanned in three consecutive blocks: immediately before, during, and immediately after stimulation. NITESGON was applied at a constant current of 1.5 mA for 20 minutes via electrodes placed over the left and right C2 nerve dermatome.

The regional amplitude of low frequency fluctuations was inspected to verify if NITESGON evoked activity changes in the LC, VTA and hippocampus. Our findings showed a significant effect for the LC (*F* = 4.34, *p* = .023), VTA (*F* = 3.42, *p* = .047) and hippocampus (*F* = 3.57, *p* = .042) when comparing the active and control groups (see fig. 6a-c). For both the LC and hippocampus, a significant increase was obtained during (LC:13.18 ±4.18 vs. 8.77 ±2.88; *F* = 11.30, *p* =.002; hippocampus: 6.30 ±3.66 vs. 4.50 ±1.31; *p* =.045) and after (LC: 13.78 ±6.21 vs. 7.71 ±2.79; *p* <.001; hippocampus: 7.35 ±4.88 vs. 4.50 ±1.44; *p* =.031) stimulation for the active group in comparison to the sham group. Before stimulation, no significant difference was obtained between the active and sham groups (LC: 9.40 ±4.50 vs. 8.98 ±1.92; *p* =.76; hippocampus: 5.26 ±2.20 vs. 5.50 ±1.64; *p* =.75). For the VTA, a significant increase was obtained during (20.01 ±5.90 vs. 14.12 ±5.34; *p* =.008) stimulation for the active group in comparison to the sham group. Before (14.96 ±4.50 vs. 14.96 ±1.92; *p* =.99) or after (15.33 ±4.98 vs. 15.46 ±13.86; *p* =.97) stimulation, no significant difference was obtained between the active and sham groups.

**Figure 6.**
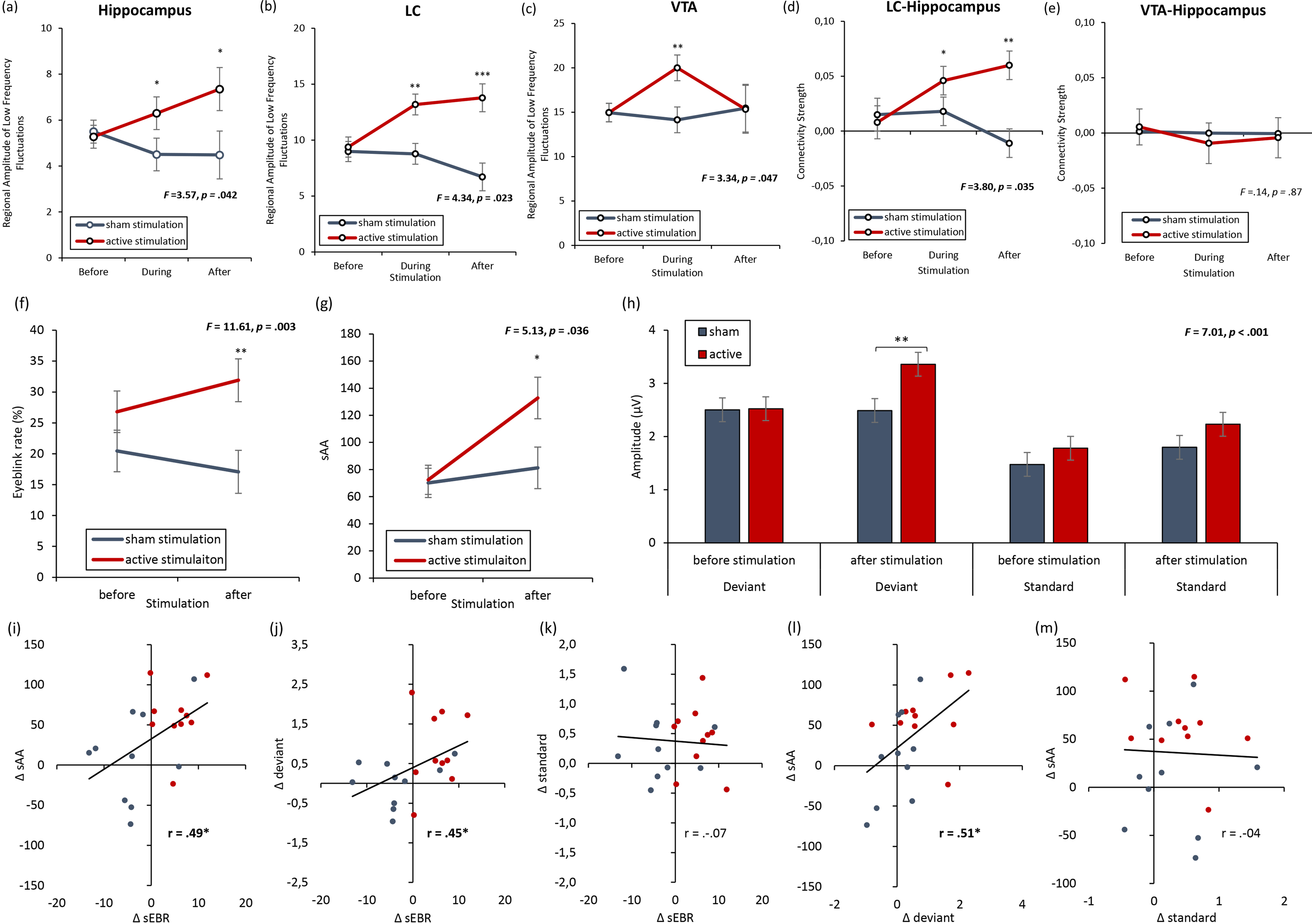
rsFMRI - Locus coeruleus and dopamine. (a,b) The locus coeruleus and hippocampus revealed increased activity during stimulation as well as after stimulation for the active NITESGON group in comparison to the sham NITESGON group. (c) The ventral tegmental area revealed increased activity during stimulation, but not after stimulation for the active NITESGON group in comparison to the sham NITESGON group. (d) Increased connectivity between the locus coeruleus and hippocampus was observed during and after stimulation for the active NITESGON group in comparison to the sham NITESGON group. (e) No significant difference in connectivity between the ventral tegmental area and hippocampus was observed when comparing the active and sham NITEGSON groups during or after stimulation. (f,g) A significant increase in spontaneous eye blink rate and sAA was observed after active NITESGON in comparison to sham NITESGON. (h) A significant increase in peak-to-peak amplitude over the left parietal electrode side was observed for the active group in comparison to the sham group for the deviant after stimulation. (i,j) A positive correlation was observed between the difference (post-pre) in spontaneous eye blink rate and the difference in sAA as well as between the difference in spontaneous eye blink rate and the difference in peak-to-peak amplitude for the deviant. (k) No significant correlation was observed between the difference in spontaneous eye blink rate and the difference in peak-to-peak amplitude for the standard. (l) A positive correlation was observed between the difference in sAA and the difference in peak-to-peak amplitude for the deviant. (m) No correlation was observed between the difference in sAA and the difference in peak-to-peak amplitude for the standard. Error bars, s.e.m. Asterisks represent significant differences (* *p* < .05; ** *p* < .01).

Furthermore, a ROI-to-ROI analysis demonstrated an effect between the right hippocampus and LC (*F* = 3.67, *p* = .039), but not between the right hippocampus and VTA (*F* = .27, *p* = .76) (see fig. 1d-e). Additionally, an increase in LC connectivity strength with the right hippocampus was seen for the active group relative to the sham group during (.052 ±.03 vs. .018 ±.06; *F* = 4.34, *p* =.047) and after stimulation (.06 ±.05 vs. -.011 ±.05; *F* = 15.25, *p* =.001). However, no significant effect was obtained between the LC and right hippocampus for the active group relative to the sham group before stimulation (.008 ±.08 vs. .015 ±.03; *F* = .09, *p* =.76).

### Experiment 7. LC-related dopamine

The previous experiment revealed activity changes in both the LC and hippocampus as well as increased connectivity between the LC and hippocampus both during and after NITESGON. Conversely, the VTA did not show changes in activity after stimulation or connectivity changes between the VTA and hippocampus during or after stimulation. However, activity changes in the VTA during NITEGSON were detected. Previous animal research has identified selective neuronal connections between the LC and VTA, implying an interaction between the LC and VTA during NITESGON may exist^21^.

A key neuromodulator in memory consolidation is dopamine (DA)^17^. DA affects plasticity, synaptic transmission and network activity in the hippocampus, and plays a critical role for hippocampal-dependent mnemonic processes by selectively enhancing consolidation of memory information^36^. Recent literature suggests a direct link between DA and the synaptic tag and capture hypothesis − the mechanism underlying behavioral tagging^37^. The core of the synaptic tag and capture hypothesis indicates that memory encoding creates the potential of long-term memory by creating a tag to be captured at a later stage (i.e., during memory consolidation) by protein synthesis dependent LTP. Suggestions are made that the signal transduction processes catalyzing this synthesis of plasticity-related proteins requires DA to stabilize new memories^13, 14^. Previous research has identified a DA agonists’ ability to chemically induce LTP specifically on synapses that are activated by test stimulation, but not those that are silent^38^, whereas a DA antagonist reduces the memory effect 24 hours after learning^39^, thus indicating that DA is central to the synaptic tag and capture hypothesis, and hence behavioral tagging^37^.

To regulate synaptic plasticity, the hippocampus receives dopaminergic input from the VTA and the LC^17, 21, 33, 36, 40, 41^. However, recent research revealed that mainly LC DA mediates post-encoding memory enhancement in the hippocampus, while the VTA does not respond to arousal (i.e., novelty). Animal research revealed that electrical stimulation of the LC increased DA levels and modulated hippocampal synaptic transmission^17, 18, 35^. Furthermore, animal studies identified activation of the LC via optogenetic stimulation caused more LTP-related memory consolidation 45 minutes after stimulation^17, 18, 35^. This could potentially explain why NITESGON applied while learning a memory task generated a long-term memory effect, but did not modify immediate learning^8, 24^.

A proxy for DA is spontaneous eye blink rate (sEBR), or the frequency of blinks per unit of time^42^. Pharmacological studies in animals and humans have shown that DA agonists elevate sEBR, whereas DA antagonist suppress sEBR^42–49^. Moreover, sEBR are altered in clinical conditions that are associated with dysfunctions of the dopaminergic system^50^.

SAA as well as neurophysiology (event-related potentials, ERP) are common proxies for LC-NA activity. More specifically, neurophysiology utilizes the P3b ERP, which peaks at 300–600 ms after a task-relevant stimulus^51, 52^, to indirectly measure LC-NA activity, thus presenting us with a strong cortical electrophysiological correlate of the LC-NA response^53^. Using an auditory oddball task, a standard P3b-evoking task, NITESGON increased peak and mean amplitude between 300 and 600 ms immediately after stimulation for the left parietal electrode site. Therefore, we hypothesized that NITESGON would induce an increase in LC related DA; shown via an increase in sEBR that would correlate with pupil diameter, sAA, and amplitude of the P3b after the application of NITESGON. sAA, sEBR and ERP were collected immediately before and immediately after 20 minutes of NITESGON was administered.

Results showed a significant interaction effect for sEBR by condition (*F* = 11.61, *p* = .003; see fig. 6f), indicating that the active group (31.90 ±10.90) had an increase in sEBR in comparison to a sham group (17.08 ±11.07; *F* = 9.10, *p* = .007) after NITESGON. Before NITESGON no significant difference (*F* = 1.77, *p* = .20) was observed in the active group (26.80 ±8.24) relative to the sham group (20.45 ±12.63) in sEBR. Also, a significant interaction effect for sAA by condition (*F* = 5,13, *p* = .036; see fig. 6g) was obtained, revealing a significant increase in sAA (*F* = 5.67, *p* = .028) after stimulation for the active group (132.82 ±51.23) in comparison to the sham group (81.22 ±45.43). No significant difference (*F* = .023, *p* = .88) was obtained when comparing the active group (72.42 ±43.77) versus the sham group (70.10 ±19.93) before stimulation. Peak-to-peak amplitude analysis for P3 electrode further showed a significant effect (*F* = 7.01, p < .001; see fig. 6h). An effect was revealed between active NITESGON and sham NITESGON after stimulation (*t* = 2.64, p = .010). In addition, a significant effect was shown for active NITESGON after stimulation in comparison to before stimulation (*t* = 2.75, p = .007). A positive correlation was obtained between the difference in sEBR and sAA (r = .49, *p* .029; see fig. 6i), peak-to-peak amplitude for deviant (r = .45, *p* .048; see fig. 6j), and peak-to-peak amplitude for standard (r = -.07, *p* = .76; see fig. 6k), respectively after NITESGON relative to before. Also, a significant correlation was obtained between sAA and peak-to-peak amplitude for deviant (r = .51, *p* .022; see fig. 6l), but not with peak-to-peak amplitude for standard (r = -.04, *p* = .87; see fig. 6m).

### Experiment 8. Dopamine

Seeing that previous research identifies DA’s vital role in memory consolidation, and NITESGON generates its effect during memory consolidation, it would be expected that blocking the DA receptor with a DA antagonist would have a direct impact on memory consolidation. To test this hypothesis, and confirm previous findings, Experiment 8 conducted a memory test 3 to 4 days after initial learning of the word-association task. We used the same setup as Experiment 1, whereby NITESGON was applied immediately after learning the word association task during visit 1.

No significant effect (*F* = .04, *p* = .85; see fig. 7a) was obtained between the participants who took a DA antagonist (71.60 ±9.18) and those who did not take a DA antagonist (70.70 ±12.04) during the learning phase on visit 1. 7 days after learning the word-associations, participants who took a DA antagonist (20.53 ±13.32) performed worse on correctly recalling words in comparison to participants who did not take a DA antagonist (40.12 ±23.68) (*F* = 5.20, *p* = .035; see fig. 7b).

**Figure 7.**
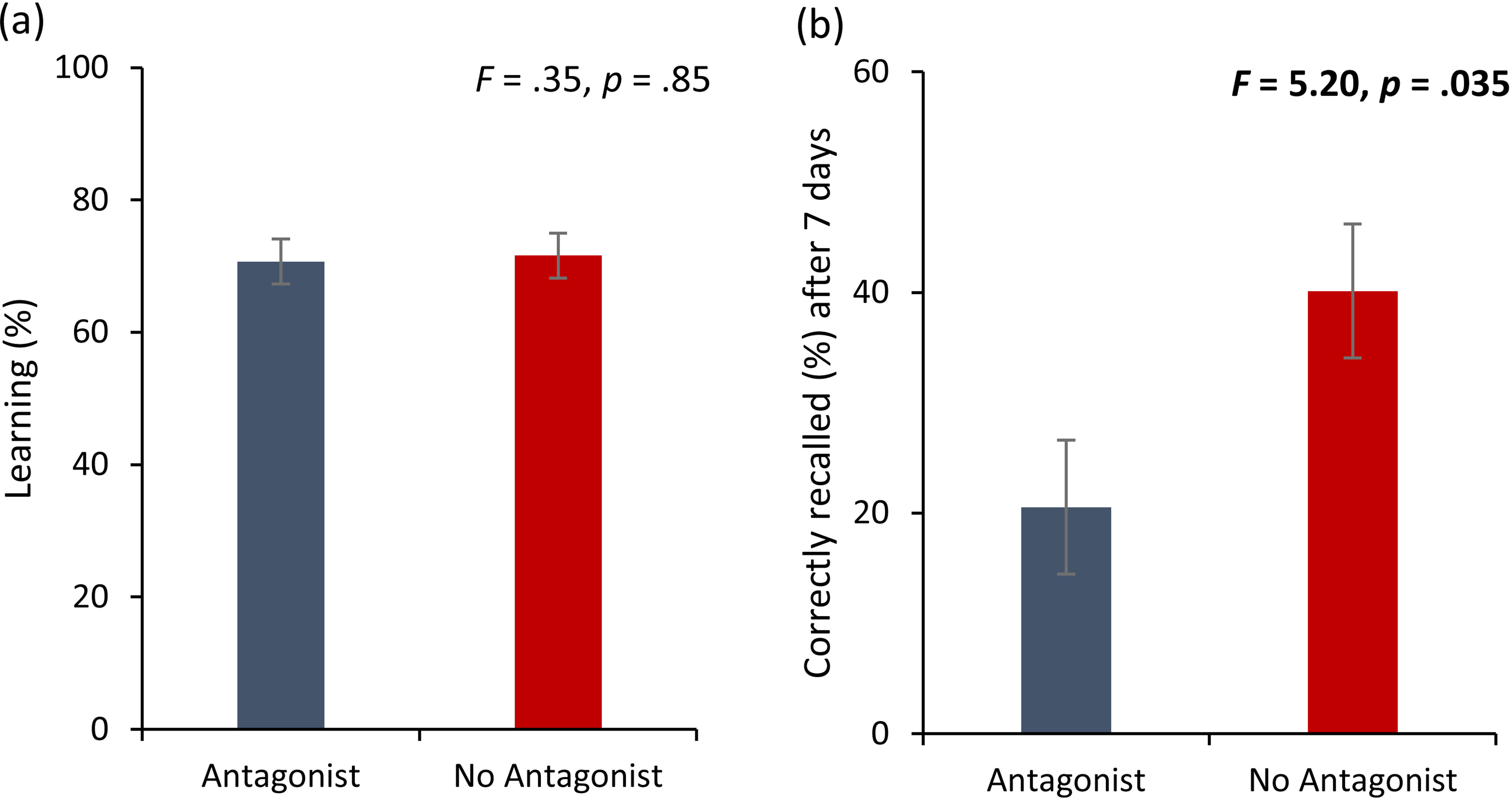
(a) No difference was observed in the cumulative learning rate after NITESGON for participants who were taking a DA antagonist in comparison to participants who were not taking a DA antagonist. (b) A significant difference was observed in the number of recalled words after 3 or 4 days for participants who were taking a DA antagonist in comparison to participants who were not taking a DA antagonist. Error bars, s.e.m. Asterisks represent significant differences (* *p* < .05; ** *p* < .01).

#### Blinding

For Experiments 1 – 7, our findings demonstrated that participants were unable to accurately determine if they were assigned to the active or sham NITESGON group, suggesting that our sham protocol is reliable and well-blinded (see fig. 8).

**Figure 8.**
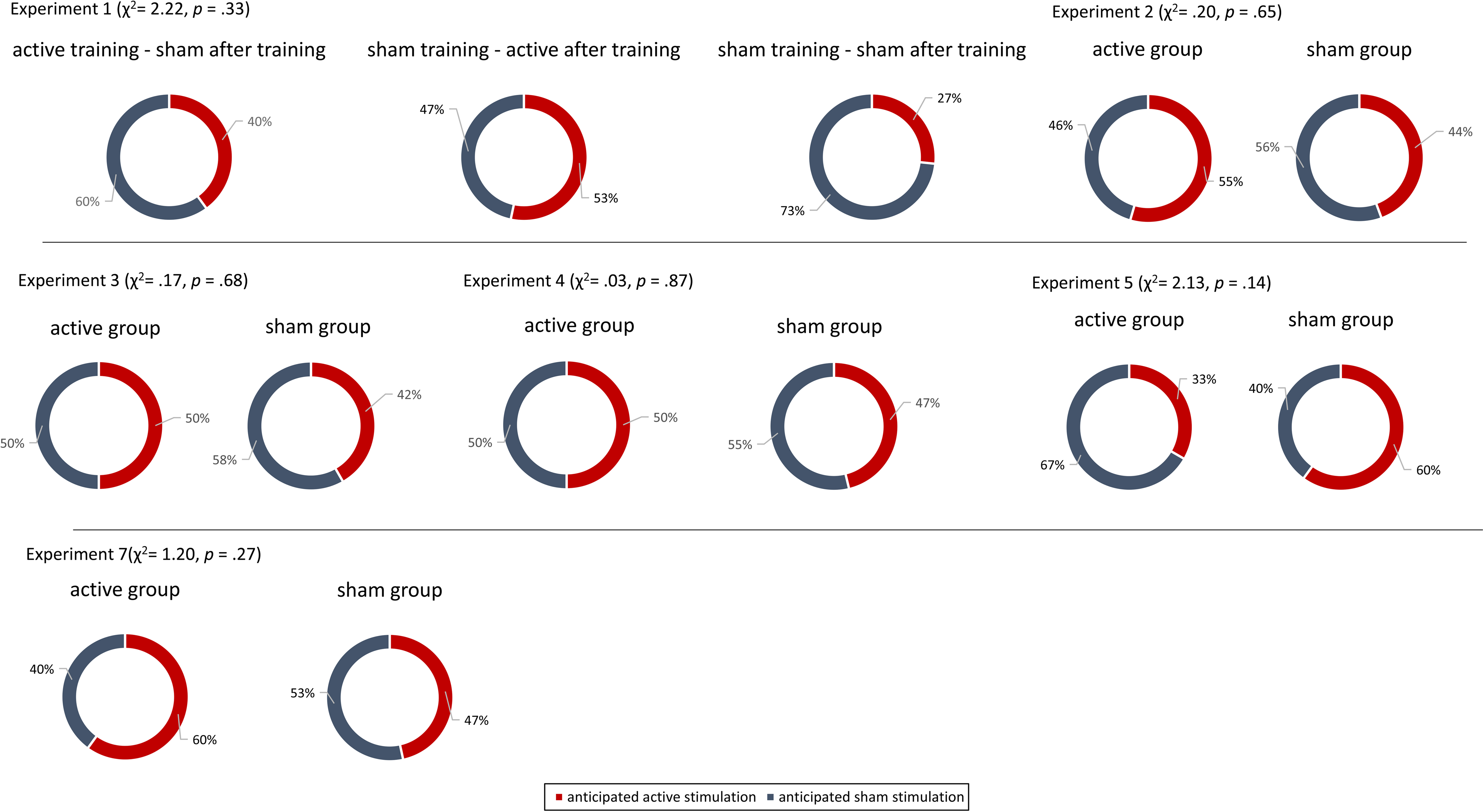
Blinding experiments. For Experiments 1 – 7, no difference was observed between the active and sham groups’ anticipation of receiving active or sham stimulation.

## Discussion

Taken together, our experiments support the hypothesis that NITESGON targeting the LC strengthens hippocampal memories via the behavioral tagging mechanism. NITESGON increases activity in the LC and hippocampus during and immediately after stimulation and increases connectivity between these two areas, thus instigating initial memory consolidation and increasing the retention of memories that are formed within a window of opportunity spanning before and after LC activation. This is in accordance with the construct of behavioral tagging, which explains how memories that would normally be forgotten will endure in memory preceding, during and following activation of the LC-pathway^18–20^.

Notably, NITESGON does not generate an immediate memory effect during learning, however, a favorable behavioral effect is seen 7 days after initial learning. Although previous research has suggested that millisecond pairing between nerve stimulation and auditory or motor learning is essential to induce targeted plasticity^5, 54, 55^, our data revealed that this design may not be as crucial as previously thought. This is comparable to the concept of behavioral tagging which suggests that there is a critical time window before and after training to transform a weak memory into a strong memory^56^. Our findings revealed that NITESGON induced a proactive and retroactive memory effect. More intriguingly, when we introduced a second word-association task (e.g.., Japanese-English) that interfered with another word-association task (e.g.., Swahili-English), we found that NITESGON diminished the interference effect. This effect induced by NITESGON did not appear to be task-specific given that we saw an advantageous effect for both spatial-navigation as well as word-association tasks. This suggests a generalized memory enhancement effect similar to prior studies of post-encoding increases in consolidation via the LC due to inducing stress and arousal^19, 20, 57^.

Previous research demonstrated that LC discharge enhances synchronization of gamma activity in the hippocampus in rats^25^ and that gamma oscillations play an important role in long-term memories and could potentially predict subsequent recall^26, 27^. Our results confirm this by revealing that NITESGON induces gamma changes in the medial temporal lobe that correlate with successful recall.

The role of the LC-NA system in synaptic plasticity and molecular memory consolidation has been well established over the past decades^21, 58^. However, recent animal studies on enhancement of memory persistence have found that LC tyrosine-hydroxylase neurons, originally defined by their canonical NA signaling, mediate post-encoding enhancement of memory in a manner consistent with possible corelease of DA from the LC axons in the hippocampus^17, 33, 34^. Interestingly, electrical stimulation of the LC increases DA levels that modulate hippocampal synaptic transmission up to 30 minutes after encoding^17, 18, 35^. Our results corroborate these findings, indicating that utilizing a DA antagonist can reduce the effect of NITESGON and that sEBR, a proxy for DA, increases after NITESGON. In addition, we demonstrated changes in sAA immediately after NTESGON that correlate with memory recall 7 days later. Previous research indicated that α-amylase is a marker of endogenous NA activity, with human fMRI showing LC activity rising concomitantly with sAA levels during the viewing of emotionally arousing slides^59^. However, original research on amylase secretion indicates that α-amylase is mediated by both DA in addition to NA^60^.

A prevailing hypothesis is that hippocampus-dependent memory is mediated by a subiculum-accumbens-pallidum-ventral tegmental area pathway via DA^61^. Our results indicated that the VTA was activated during NITESGON, however, the VTA activation ceased post stimulation. No increased connectivity was revealed between the VTA and hippocampus during or after NITESGON. This corresponds with recent research that suggests hippocampal projections in the VTA are sparse^17, 33, 34^ and therefore may only have a limited role in late-phase LTP^17, 33, 34^. However, it is possible that the VTA indirectly contributes to the formation of memories via other brain areas. Recent animal research has suggested that VTA DA neurons project to the amygdala and may contribute to fear memory in addition to the LC^62, 63^, and that the VTA may contribute to synaptic consolidation independently and complementary to the LC^23^. Further experimental investigations are needed in order to establish this.

Although our results reflect the putative tag and capture mechanism, future research needs to be conducted to determine whether such a mechanism explains our behavioral effects shown here. The theory that the effect of NITESGON is generated during memory consolidation is supported by the observation that effects were not seen immediately but were demonstrated 7 days later and were not affected by sleep. This is analogous with studies showing that arousal-mediated consolidation effects are time-dependent, but less dependent on sleep^32^.

In conclusion, our work provides evidence that NTESGON is involved in the consolidation of information rather than encoding. Our findings support an implication previously put forward in the formulation of the synaptic tag and capture mechanism proposing late-phase LTP of synaptic activity could explain enhanced memories. As deficits in episodic memory specifically related to memory consolidation are one of the earliest detectable cognitive abnormalities in amnestic mild cognitive impairment and Alzheimer’s disease^64–66^, NITESGON might have the potential of improving memory recall by hampering the disruption of memory consolidation.

## Methods

All experiments were designed as a prospective, double-blinded, placebo-controlled, randomized parallel-group study where the researcher and the participant were blinded to the stimulation conditions. Experiment 6 was alone a single-blinded study where the participant was blinded to the stimulation condition, but not the researcher. All experiments were in accordance with the ethical standards of the Declaration of Helsinki (1964). Experiments 1-7 were approved by the Institutional Review Board at the University of Texas at Dallas. All participants signed a written informed consent.

In all experiments, direct current was transmitted via a saline-soaked (1.3% saline) pair of synthetic sponges (5 cm x 7 cm) and was delivered by a specially developed, battery-driven, constant current stimulator with a maximum output of 10 mA (Eldith^©^; http://www.neuroconn.de). For each participant receiving NITESGON, the anodal electrode was placed over the left C2 nerve dermatome and the cathodal electrode was placed over the right C2 dermatome. To maintain consistency across all participants, research assistants were trained to map out the placement according to the length of the participant’s head.

To minimize skin sensations and to acclimate participants to the stimulation types, the current intensity was ramped-up (gradually increasing) until it reached its programmed maximum output (1.5 mA). After stimulating for the desired duration per the group (active or sham), the current was ramped-down (gradually decreased) denoting the end of the stimulation. The impedance under each electrode was maintained under 10 kΩ. The ramp-up, ramp-down and stimulation times were different depending on condition (active vs sham) and experimental needs.

### Experiment 1: NITESGON during or immediately after training

#### Participants

Participants were 48 healthy, right-handed, native-English speaking adults (24 males, 24 females; mean age was 20.02 years, *Sd* = 1.75 years) with a similar educational background (i.e., enrolled as undergraduate students at UT Dallas) with normal to corrected vision, who all had the maximum score on the Mini Mental State Examination. Participants were screened (e.g., tES contraindications, neurological impairments, not participated in a tES study) prior to enrolling into the study. None of the participants had a history of major psychiatric or neurological disorders, or any tES contraindications, including previous history of brain injuries or epileptic insults, cardiovascular abnormalities, implanted devices, taking neuropsychiatric medications, prescribed stimulants use, or chronic use of illicit drugs (i.e., marijuana and cocaine).

Participants were excluded from the study if screening discovered they were familiar with Swahili/Arabic language or Swahili culture due to the nature of the stimuli. Furthermore, participants received instructions advising them to abstain from the following products for the associated time window prior to their study session: dental work for 48-hours, alcohol for 24-hours, caffeine and nicotine for 16-hours, and hair styling products the day of. Participants provided written, informed consent on the day of the study session.

#### Word-association task

Associative memory performance was measured using a computerized Swahili-English verbal paired-associative learning task. This task was adapted from a well-established study design published in Science by Karpicke and Roediger^67^. Using a SDT_N_ paradigm (S: study phase, D: distraction phase, T_N_: test phase with non-recalled word pairs), participants were instructed to read and remember 75-sequentially presented Swahili-English (e.g., Swahili: bustani, English: garden) word pairs made up of common day-to-day words. The Swahili-English word pairs were taken from the study by Nelson and Dunlosky^68^. Participants had the opportunity to learn the list of 75 word pairs repetitively across a total of four alternating study and test periods each. During the study period, the word pairs were presented together on a computer screen for 5 seconds with the Swahili word on top and the English translation at the bottom (5 x 75 = 375 sec). The study period was followed by a cued-recall test-period: Swahili cue words were presented for 8 seconds each during which participants had to type-in the correct English translation remembered from the study period. Correctly recalled word pairs were dropped from further testing but remained to be studied in each subsequent learning period (i.e., 4 blocks of studying 75 word pairs). The order of the words being studied or tested were randomized. Previous research has demonstrated the critical role of retrieval practice in learning of a new foreign language; therefore, the paradigm ensures that all participants were well exposed to the stimuli and avoided a ceiling effect^67^.

#### NITESGON

There were three groups – active NITESGON during learning (i.e., study phases of the word association task) and sham NITESGON immediately after the word-association task; sham NITESGON during learning and active NITESGON immediately the word-association task in the memory consolidation period and sham NITESGON both during and after learning of the word-association task – with 16 participants each. Active NITESGON consisted of ramp-up time of 30 seconds followed by a constant current of 1.5 mA (current density 42.28 μA/cm2) during each of the 4 study blocks, resulting in a total stimulation time of 25 minutes (i.e., 375 seconds × 4 blocks) and ramp-down time of 30 seconds. For the sham NITESGON group, the current intensity was ramped-up to 1.5 mA over 30 seconds and immediately ramped-down over 30 seconds. Hence, sham NITESGON only lasted 60 seconds per study period, resulting in a total time of 240 seconds (60 seconds x 4 blocks) of stimulation when delivered during the study phase. For the group that received active NITESGON after the word-association task, this consisted of 30 seconds ramp-up and ramp-down time with 25 minutes of constant current stimulation at 1.5 mA. The sham NITESGON delivered after the word-association task only consisted of 30 seconds of ramp-up and ramp-down time resulting in 60 seconds of stimulation. The rationale behind the sham procedure was to mimic the transient skin sensation at the beginning of active NITESGON without producing any conditioning effects on the brain.

#### Resting-state EEG

Continuous EEG data was collected from each participant pre- and post-NITESGON procedures. The data was collected using a 64 channel Neuroscan Synamps2 Quick Cap configured per the International 10-20 placement system with the midline reference located at the vertex and the ground electrode located at AFZ using the Neuroscan Scan 4.5 software (Neuroscan, http://compumedicsneuroscan.com). The impedance on each electrode was maintained at less than 5 kΩ. The data were sampled using the Neuroscan Synamps2 amplifier at 500 Hz with online band-pass filtering at .1–100 Hz.

Eyes-closed recordings (sampling rate = 1 kHz, band passed DC–200 Hz) were obtained in a dark room which was dimly lit with a small lamp with each participant sitting upright in a comfortable chair; data collection lasted approximately 5 minutes. Participants were instructed not to drink alcohol 24-hours prior to EEG recording or caffeinated beverages one hour before recording to avoid alcohol- or caffeine-induced changes in the EEG stream. The alertness of participants was checked by monitoring both slowing of the alpha rhythm and appearance of spindles in the EEG stream to prevent possible enhancement of the theta power due to drowsiness during recording^69^. No participants included in the current study showed such EEG changes during measurements.

#### Saliva Collection

Saliva was collected twice during each experiment: once immediately prior to NITESGON stimulation and once immediately after NITESGON stimulation. When the participants were ready to collect saliva, they were requested to gently tip their head backwards and collect saliva on the floor of their mouth and when ready, passively drool into the collection aid mouthpiece provided by Salimetrics laboratory (Salimetrics, LLC, USA; https://salimetrics.com). The participants were requested to collect 2 ml of saliva in one straight flow and avoid breaks between drool as much as possible. The length of time to collect 2 mL of saliva was noted and the timer was started only when participants began to passively drool into the tube. All saliva samples were stored in 2 mL cryovials, and immediately stored in an −80° C laboratory freezer. Prior to saliva collections, all participants were instructed to avoid food, sugary drinks, excess caffeine, nicotine, and acidic drinks for at least one hour before collecting the saliva samples. Participants were also instructed to avoid alcohol and vigorous exercise 24-hours prior to and avoid dental work 48-hours prior to their appointment. In addition, participants were instructed not to brush their teeth within 45-minutes of sample collection in order to avoid any risk of lowering pH levels and influencing bacterial growth. If the study was scheduled for the afternoon, participants were requested to avoid taking naps during the day. Upon completion of the collection procedures, all saliva samples were packed in dry ice and sent to the Salimetrics laboratory for analysis. We chose to use salivary α-amylase (sAA) as a biomarker for norepinephrine as it provided a noninvasive yet valid indicator of central sympathetic nervous system activation^70^. SAA levels have been shown to co-vary significantly with circulating NA levels, with human fMRI showing locus coeruleus activity rising simultaneously with sAA levels during viewing of emotionally arousing slides^59^.

#### Procedure

Eligible participants were scheduled for two visits to complete the study. Visit 1 consisted of the word-association task and administration of NITESGON. Participants were randomly assigned to one of three groups during the study period. The researcher who controlled the NITESGON device was not involved in instructing the participant; this was performed by a second researcher who was blind to the stimulation protocol. Resting-state EEG (rsEEG) and saliva were collected twice for all participants, once immediately before and once immediately after NITESGON application. Participants were asked to refrain from studying or searching for the learned word pairs throughout the week. Participants returned 7 days after their first visit for memory testing to measure possible (long-term) effects on associative memory performance, but did not receive NITESGON, nor were they able to review word-pairs. A third researcher who was not responsible for the task or NITESGON conducted the second visit.

#### EEG preprocessing

For the EEG preprocessing, the data was resampled to 128 Hz, band-pass filtered (Finite Impulse Response filter) to 2–44 Hz, and re-referenced to the average reference using EEGLAB 14_1_1b^71^. The EEG data was then plotted for a careful inspection for artifacts. All episodic artifacts suggestive of eye blinks, eye movements, jaw tension, teeth clenching, or body movements were manually removed from the EEG stream. In addition, an independent component analysis (ICA) was conducted to further verify whether all artifacts were excluded.

#### EEG source localization

Standardized low-resolution brain electromagnetic tomography (sLORETA) was used to estimate the intracerebral electrical sources that generated the scalp-recorded activity in each of the gamma frequency bands (30.5– 44 Hz)^72^. sLORETA computes neuronal activity as current density (A/m^2^) without assuming a predefined number of active sources. The sLORETA solution space consists of 6,239 voxels (voxel size: 5 × 5 × 5 mm) and is restricted to cortical gray matter and hippocampi, as defined by the digitized Montreal Neurological Institute (MNI) 152 template^73^. Scalp electrode coordinates on the MNI brain are derived from the international 10–20 system^74^.

The tomography of sLORETA has received considerable validation from studies combining sLORETA with other more established spatial localization methods such as fMRI^75, 76^, structural MRI^77^, and PET^78–80^. Further sLORETA validation is based on accepting as ground truth that the localization findings obtained from invasive, implanted depth electrodes, of which there are several studies in epilepsy^81, 82^ and cognitive ERPs^83^.

#### Statistics task

For visit 1 learning, a one-way ANOVA was conducted with the cumulative learning rate over the different study periods as the dependent variable and three groups as the between-subjects variable. To look at the memory effect (recall) 7 days after learning, we applied a one-way ANOVA with the group as the between-subjects variable and correctly recalled words as the dependent variable.

#### Statistics saliva

Using the saliva collected via the passive drool method, sAA levels were measured. We conducted a repeated measures ANOVA with groups as between-subjects variable and sAA as within-subjects variable. A simple contrast analysis was applied to compare the different conditions using a Bonferroni correction.

#### Statistics EEG – whole brain analysis

A whole brain analysis was used to compare gamma activity before and after NITESGON. These activity changes were then correlated with the number of words recalled during visit 2, 7 days after the learning task, using a Pearson correlation. Non-parametric statistical analyses of functional sLORETA images (statistical nonparametric mapping) were performed for each contrast employing a t-statistic for paired groups and corrected for multiple comparisons (P < 0.05). The significance threshold for all tests was based on a permutation test with 5000 permutations and corrected for multiple comparisons^84^.

### Experiment 2 – NITESGON during second task – retroactive strengthening of memories

#### Participants

Participants were 20 healthy, right-handed, native-English speaking adults (9 males, 11 females; mean age was 21.11 years, *Sd* = 2.03 years) with a similar educational background (i.e., enrolled as undergraduate students at UT Dallas). Participants were screened and enrolled similar to Experiment 1.

#### Word-association task (task 1)

Associative memory performance was measured using the same computerized Swahili-English verbal paired-associative learning task used in Experiment 1, however, the task consisted of 3 study periods in which participants were asked to read and remember 50 Swahili-English word pairs in each study period (50 x 5 = 250 seconds).

#### Object-Location task (task 2)

Participants partook in a second memory performance task immediately after the word-association task. The second memory task consisted of a spatial navigation object-location association task that was based on previous research^85^. Using the same SDT_N_ paradigm, participants were instructed to view and remember 50 sequentially presented objects locations on a blue-red-gray background grid with an eye-to-screen distance of ∼24 inches across three study-test blocks. The objects consisted of black and white line drawings from the Boston Naming Test; 10 objects from each of the following categories were used: animals, foods, modes of transportation, tools, and household objects. Each image appeared in a randomized order at a randomized location. Objects were presented one at a time for 3000 ms each (1000-ms ISI). Objects were presented within a white-box background (4.88 x 4.88 cm) and had a red dot superimposed at the object center to mark the precise location. Participants were instructed to study and remember the object-locations as accurately and precisely as possible. After each study phase, a cued-recall test was administered. During the test period, the studied objects were presented one at a time in the center of the screen (in a randomized order), and participants were required to recall the studied locations. At the beginning of every trial, a 2000 ms fixation cross at the center of the screen was presented. After this 2000 ms period, participants were able to use the mouse to move the object from the center of the screen to its recalled location and click a button on the mouse to indicate its final location.

The Swahili-English verbal associative task was used as task 1 and the spatial navigation object-location task was used as task 2 for all participants.

#### NITESGON

All participants received sham NITESGON during each study period of task 1 using the following parameters: a 5 second ramp up period, followed by a constant current of 1.5 mA for 15 seconds, ending with a ramp down period of 5 seconds, allowing for an emulated sensation of the active NITESGON. For task 2, 10 participants received active NITESGON and 10 participants received sham NITESGON. Sham stimulation parameters were the same as used in task 1 and stayed consistent in each of the three study periods of task 2. Participants given active NITESGON received a 5 second ramp-up period, followed by a constant current of 1.5 mA for 250 seconds, and finished with a 5 second ramp down period during each of the 3 study periods of task 2 on the first day. Thus, the total simulation time for the active group was 750 seconds (i.e., 250 seconds × 3 study periods) and the sham group was 45 seconds (i.e., 15 seconds × 3 study periods). Just before the first study period of the first task participants NITESGON was delivered for 15 seconds to help participants habituate to the sensation and to check if they were comfortable with the sensation.

#### Resting-state EEG

Continuous EEG data was collected from each participant pre- and post-NITESGON procedures as detailed in experiment 1.

#### Saliva collection

Saliva was collected twice during each experiment: once immediately prior to NITESGON stimulation and once immediately after NITESGON stimulation as detailed in experiment 1.

#### Procedure

Eligible participants were scheduled for two visits to complete the study. Visit 1 consisted of the word-association task (i.e., task 1) paired with sham stimulation, and then were randomly assigned to either the active or sham NITESGON condition for the spatial navigation task (i.e., task 2). The researcher who controlled the NITESGON device was not involved in instructing the participant; this was performed by a second researcher who was blind to the stimulation protocol. rsEEG and saliva were collected twice for all participants, once immediately before and once immediately after NITESGON application. Participants were asked to refrain from studying or searching for the learned word pairs throughout the week. Participants returned 7-days after their first visit for memory testing on both task 1 and task 2 to measure possible (long-term) effects on associative memory performance, but did not receive NITESGON, nor were they able to review word-pairs or objects’ locations. A third researcher who was not responsible for the task or NITESGON conducted the second visit.

#### EEG preprocessing and source localization

The continuous EEG data was preprocessed and the source-level gamma activity pre- and post-NITESGON procedures for the two groups was determined as detailed in experiment 1.

#### Statistics task

For visit 1 learning, a MANOVA was conducted with the cumulative learning rate over the different study periods for both tasks as the dependent variable and group as the between-subjects variable. To look at the memory effect (recall) 7 days after learning, we applied a MANOVA with groups as the between-subjects variable and correctly recalled words on both tasks as the dependent variable. For both analyses, if significant, two separate one-way ANOVAs were applied with groups as the between-subjects variable and correctly recalled words as dependent variable for task 1 or task 2 respectively.

#### Statistics saliva

Using the saliva collected via the passive drool method, sAA levels were measured which were compared between the groups as detailed in experiment 1.

#### Statistics EEG – whole brain analysis

A whole brain analysis was used to compare gamma activity before and after NITESGON. This activity was correlated with the number of correctly recalled items (words/locations) 7 days later as detailed in experiment 1.

### Experiment 3 - NITESGON during first task - proactive strengthening of memories

#### Participants

Participants were 24 healthy, right-handed, native-English speaking adults (13 males, 11 females; mean age was 20.83 years, *Sd* = 2.21 years) with a similar educational background (i.e., enrolled as undergraduate students at UT Dallas). Participants were screened and enrolled similar to Experiment 1.

#### Word-association task (task 1)

Associative memory performance was measured using the same computerized Swahili-English verbal paired-associative learning task used in Experiment 2.

#### Object-location task (task 2)

Participants partook in a second memory performance task immediately after the word-association task consisting of a spatial navigation object-location association task used in Experiment 2.

#### NITESGON

The same device, electrode placement, and active and sham NITESGON parameters described in Experiment 2 were used. Differing from Experiment 2, Experiment 3 had all participants receive sham NITESGON during each study period of task 2 as opposed to task 1. 12 participants received active NITESGON and 12 participants received sham NITESGON during the second task.

#### Resting-state EEG

Continuous EEG data was collected from each participant pre- and post-NITESGON procedures as detailed in experiment 1.

#### Saliva collection

Saliva was collected twice during each experiment: once immediately prior to NITESGON stimulation and once immediately after NITESGON stimulation as detailed in experiment 1.

#### Procedure

Eligible participants were scheduled for two visits to complete the study. Visit 1 consisted of the word-association task (i.e., task 1) paired with either active or sham NITESGON, and spatial navigation task (i.e., task 2) paired with sham NITESGON. The researcher who controlled the NITESGON device was not involved in instructing the participant; this was performed by a second researcher who was blind to the stimulation protocol. rsEEG and saliva were collected twice for all participants, once immediately before and once immediately after NITESGON application. Participants were asked to refrain from studying or searching for the learned word pairs throughout the week. Participants returned 7-days after their first visit for memory testing on both task 1 and task 2 to measure possible (long-term) effects on associative memory performance, but did not receive NITESGON, nor were they able to review word-pairs or objects’ locations. A third researcher who was not responsible for the task or NITESGON conducted the second visit.

#### EEG preprocessing and source localization

The continuous EEG data was preprocessed and the source-level gamma activity pre- and post-NITESGON procedures for the two groups was determined as detailed in experiment 1.

#### Statistics task

The learning in visit 1 and memory performance in visit 2 was compared between the groups as detailed in experiment 2.

#### Statistics saliva

Using the saliva collected via the passive drool method, sAA levels were measured which were compared between the groups as detailed in experiment 1.

#### Statistics EEG – whole brain analysis

A whole brain analysis was used to compare gamma activity before and after NITESGON. This activity was correlated with the number of correctly recalled items (words/locations) 7 days later as detailed in experiment 1.

### Experiment 4 – NITESGON during interference task

#### Participants

Participants were 31 healthy, right-handed, native-English speaking adults (15 males, 16 females; mean age was 21.36 years, *Sd* = 2.42 years) with a similar educational background (i.e., enrolled as undergraduate students at UT Dallas). Participants were screened and enrolled similar to Experiment 1. Experiment 4 added familiarity of Japanese language or culture to the participant screening procedure, if indicated, the participant was excluded from the study due to the nature of the stimuli.

#### Word-association tasks

Associative memory performance was measured using two computerized verbal paired-associate learning tasks. Similar to Experiment 2 and 3, one task comprised of the Swahili-English vocabulary learning, and the second task consisted of a newly introduced Japanese-English (e.g., Japanese: Kumo, English: cloud) word-association task. The Japanese-English word-association task used the same Swahili-English word-pairs, however, the Swahili words were replaced by Japanese words.

#### NITESGON

The same device, electrode placement, and active and sham NITESGON parameters described in Experiment 2 was used. 16 participants received active NITESGON and 16 participants received sham NITESGON during the first task, where everyone received sham NITESGON during the second task.

#### Resting-state EEG

Continuous EEG data was collected from each participant pre- and post-NITESGON procedures as detailed in experiment 1.

#### Saliva collection

Saliva was collected twice during each experiment: once immediately prior to NITESGON stimulation and once immediately after NITESGON stimulation as detailed in experiment 1.

#### Procedure

Eligible participants were scheduled for two visits to complete the study. Visit 1 consisted of two word-association tasks, whereby task 1was paired with either active or sham NITESGON followed by a second word-association task (i.e., task 2) paired with sham NITESGON. The order of the two word-association tasks was randomized over the participants in a 1:1 ratio to remove a possible order effect. The researcher who controlled the NITESGON device was not involved in instructing the participant; this was performed by a second researcher who was blind to the stimulation protocol. rsEEG and saliva were collected twice for all participants, once immediately before and once immediately after NITESGON application. Participants were asked to refrain from studying or searching for the learned word pairs throughout the week. Participants returned 7-days after their first visit for memory testing on both task 1 and task 2 to measure possible (long-term) effects on associative memory performance, but did not receive NITEGSON, nor were they able to review word-pairs. A third researcher who was not responsible for the task or NITESGON conducted the second visit. As during the first visit, the two word-association tasks were randomized over the participants in a 1:1 ratio to remove a possible order effect.

#### EEG preprocessing and source localization

The continuous EEG data was preprocessed and the source-level gamma activity pre- and post-NITESGON procedures for the two groups was determined as detailed in experiment 1.

#### Statistics task

For visit 1 learning, a repeated measures ANOVA was applied with the cumulative learning rate over the different study periods for both tasks as within-subjects variable and group (active vs. sham) as between-subjects variable. A similar analysis was applied for the memory effect (recall) 7-days after learning. A simple contrast analysis was applied to compare the difference conditions using a Bonferroni correction. In addition, an interference effect was calculated by subtracting the memory recall 7-days after learning the second task from the first task. This number give a proxy of interference. A one-way ANOVA was applied with the interference effect as dependent variable and group (active vs. sham) as between-subjects variable. Lastly, to see if the interference effect was significantly different from zero (i.e., no interference effect) for both the active and sham group, a one-sample t-test was used.

#### Statistics saliva

Using the saliva collected via the passive drool method, sAA levels were measured which were compared between the groups as detailed in experiment 1.

#### Statistics EEG – whole brain analysis

A whole brain analysis was used to compare gamma activity before and after NITESGON. This activity was correlated with the number of correctly recalled items (words/locations) 7 days later as detailed in experiment 1.

### Experiment 5 – Effect of NITESGON is not sleep dependent

#### Participants

Participants were 20 healthy, right-handed, native-English speaking adults (11 males, 9 females; mean age was 21.18 years, *Sd* = 1.951 years) with a similar educational background (i.e., enrolled as undergraduate students at UT Dallas). Participants were screened and enrolled similar to Experiment 1.

#### Word-association task

Associative memory performance was measured using the same computerized Swahili-English verbal paired-associative learning task used in Experiment 1.

#### NITESGON

All participants received active NITESGON during each of the four study periods on visit 1 using the following parameters: a 5 second ramp-up period, followed by a constant current of 1.5 mA for 375 seconds (75 word-pairs x 5 seconds), and finished with a 5 second ramp-down period. The total stimulation time was 25 minutes (i.e., 375 sec × 4 blocks). Before the start of the first study period, an additional 15 second habituation period was added to make sure the participants got used to the sensation.

#### Procedure

Eligible participants were scheduled for two visits to complete the study. Visit 1 consisted of the word-association task paired with active NITESGON. Participants were randomly assigned to one of the two groups (sleep vs no sleep). 10 participants learned the word-association task at 8:00 a.m. and were tested the same day at 8:00 p.m., while the other 10 participants learned the word-association task at 8:00 p.m. and were tested the next day at 8:00 a.m. after a night of sleep. Participants were asked to refrain from studying or searching for the learned word pairs for at least the next 12-hours. The researcher who controlled the NITESGON device was not involved in instructing the participant; this was performed by a second researcher who was blind to the stimulation protocol. A third researcher who was not responsible for the task or NITESGON conducted the second visit (12-hours later).

#### Statistics task

A one-way ANOVA with group (sleep vs no sleep) as between-subject variable and number of words correctly recalled as dependent variable was performed.

### Experiment 6 – LC – Hippocampus activity & connectivity

#### Participants

Participants were 32 healthy, right-handed, native-English speaking adults (16 males, 16 females; mean age was 25.32 years, *Sd* = 2.65 years) with a similar educational background (i.e., enrolled as undergraduate students at UT Dallas). Participants were screened and enrolled similar to Experiment 1. Experiment 6 added the exclusion of those participants who had any contraindication for MRI (i.e., metallic implants, pregnancy, claustrophobia).

#### Resting-state fMRI

The resting state fMRI data were collected on a 3T MR scanner (Achieva, Philips, Netherlands) using a 32-channel SENSE phased-array head coil. The dimension of the coil was 38 (height) × 46 (width) × 59 (length) cm^3^. During scanning, foam padding and earplugs were used to minimize the head movement and scanner noise. An MR-compatible, battery powered NITESGON system manufactured by MR NeuroConn Co. (Germany) was applied to each participant inside the MR scanner. All the operating parts and devices that go into the scanner room were MR-compatible and everything else was in the control room, connected via the waveguide. The NITESGON system was fully charged before each session and its impedance level was measured regularly to test if it was maintained at approximately 5 kΩ on each end (i.e., 10 kΩ total).

The MR session with NITESGON was divided into three consecutive blocks of scanning: before stimulation, during stimulation, and after stimulation. At the beginning of the pre-stimulation session, routine survey and T1 anatomical images were acquired for a total time of 5-minutes. Before acquiring the T1 image, saline-soaked NITESGON electrodes were positioned on the subject for three consecutive blocks of rsfMRI. For each of the scanning blocks, we acquired 20-minute long rsfMRI images.

For the T1 (MPRAGE) anatomical scan, parameters were repetition time (TR) of 2300 ms, an echo time (TE) of 2.94 ms, an inversion time (TI) of 900 ms, and a flip angle of 9◦. 160 sagittal slices were taken, using a matrix size of 256 × 256 mm^2^, at a 1 × 1 × 1 mm^3^ resolution.

Resting state fMRI sequences were acquired with the following imaging parameters (echo-planar imaging protocol): TR/TE = 3000/30 ms, FOV = 220 × 220 mm^2^, matrix = 64 × 64, number of slices = 53 with voxel size = 3 × 3 × 4 mm^3^ with no gap between slices. The total number of acquired volumes was 400, counting for 20 minutes. Preprocessing steps can be found in supplementary materials.

#### NITESGON

Shielded cables connected the MR-compatible box and electrodes, and the stimulation data was transferred via the CAT.6 LAN cable that runs throughout the MR scanner room to the non-MR-compatible stimulation devices in the control room. For the active NITESGON group the current was ramped-up for 30 seconds followed by a constant current of 1.5 mA for 20 min and a 10 second ramp-down period. For sham NITESGON, the current was ramped-up over 30 seconds to reach the intensity of 1.5 mA followed by 15 seconds of constant current stimulation at 1.5 mA and 10 seconds ramp-down. Hence, sham NITESGON only lasted 15 seconds (as opposed to 20 minutes in the active group).

#### Procedure

Participants were scanned immediately before, during, and immediately after the NITESGON stimulation. The researcher who controlled the NITESGON device was not blinded to the stimulation group but the participant was blinded to which stimulation group they were placed in. 16 participants received active NITESGON and 16 received sham NITESGON.

#### Statistics rsfMRI

A functional connectivity analysis was performed using the CONN toolbox. The regions of interest (ROI) considered in the analysis were the right hippocampus (based on previous findings^8^), LC and VTA. Both the LC and VTA were selected using probabilistic atlas (as conducted in a study across 44 adults by localizing its peak signal level in high-resolution T1 turbo spin-echo images and verified the location using post-mortem brains)^86^. The probabilistic templates were created using processing steps specifically designed to address these difficulties^86^. To remove potential artifacts such as head motion, respiration, and other global imaging artifacts including potential stimulation effects, we regressed out the global average brain signal.

We conducted a regional amplitude of low-frequency fluctuation (rALFF) analysis for the LC, VTA and hippocampus. The time series for each voxel of each ROI was transformed to the frequency domain and the power spectrum was then obtained. Since the power of a given frequency is proportional to the square of the amplitude of this frequency component, the square root was calculated at each frequency of the power spectrum and the averaged square root was obtained across 0.01–0.17 Hz at each voxel. This averaged square root was taken as the rALFF^87^. The rALFF of each voxel was divided by the individual global mean of the rALFF within a brain-mask, which was obtained by removing the tissues outside the brain using software MRIcron. Spatial smoothing was conducted on the maps with an isotropic Gaussian kernel of 8 mm of full width at half-maximum. A repeated measures ANOVA was used including group (active vs sham) as between-subjects variable and rALFF before, during and after NITESGON as within-subjects variable for the different ROIs (ALFF for the VTA, LC and hippocampus). A simple contrast analysis was included to compare the difference between active and sham stimulation for each ROI before, during, and after stimulation separately including a correction for multiple comparison (Bonferroni correction).

In addition, the average BOLD time series across all voxels within the ROIs were extracted from the smoothed functional images. A partial correlation analysis was performed, and the resulting *r*-value converted to Fisher’s *Z*-transformed coefficients were used for further statistical analyses. The *Z*-transformed connectivity weights were compared between the active and sham groups for the LC and hippocampus, LC and VTA, and VTA and hippocampus, respectively using a repeated measures ANOVA. A simple contrast analysis was applied to compare the different the active and sham condition using a Bonferroni correction.

### Experiment 7 – LC-related dopamine

#### Participants

Participants were 24 healthy, right-handed, native-English speaking adults (12 males, 12 females; mean age was 23.83 years, *Sd* = 2.88 years) with a similar educational background.. Participants were screened and enrolled similar to Experiment 1.

#### NITESGON

Active NITESGON stimulation consisted of a ramp-up period of 5 seconds, followed by constant current of 1.5 mA for 20 minutes and ramp-down period of 5 seconds. Sham NITESGON only consisted of a ramp-up period of 5 seconds to reach the intensity of 1.5 mA and an immediate ramp-down period of 5 seconds. 12 participants received active NITESGON and 12 participants received sham NITESGON.

#### Electrophysiological recordings

Continuous EEG data was collected from each participant in response to the auditory oddball paradigm, before and after the application of NITESGON. The auditory oddball task is a simple and well-established paradigm for the investigation of the robust P3b component which has a predictable standard tone and an unpredictable deviant tone^88^. The data was collected using a 64-channel Neuroscan Synamps^2^ Quick Cap configured per the International 10-20 placement system with the midline reference located at the vertex and the ground electrode located at AFZ using the Neuroscan Scan 4.5 software. The impedance on each electrode was maintained at less than 5 kΩ. The data were sampled using the Neuroscan Synamps^2^ amplifier at 500 Hz with online band-pass filtering at .1–100 Hz. Data was preprocessed using Matlab and EEGLAB in a manner similar to the original paper that showed a relationship between ERP and locus coeruleus–noradrenergic arousal function^88^.

#### Peak-to-peak P3b amplitude

Peak-to-peak amplitude was defined as the amplitude difference between the N200 peak and the P300 peak for the P3 electrode. The N200 component was identified as the most negative peak between 200 and 375 ms after the stimulus onset. The P300 component was identified as the most positive peak between 250 and 600 ms after the stimulus onset.

#### Spontaneous eyeblink rate (sEBR)

To retain the eyeblinks, the eyeblink rate was calculated using the data before cleaning the artifacts using an independent component analysis. Furthermore, the continuous dataset before epoching was used to visualize the entire temporal profile of the eyeblink potential to avoid any cutting-off of the potential due to epoching. An eyeblink was determined to be a sharp negative peak followed immediately by a positive peak located in the frontal electrodes such as FP1, FP2 and FPz. In some cases, the negative peak was not prominent, but the positive peak was a signatory. The topography of this potential was observed to have a clear dipole covering the frontal and fronto-temporal electrodes. This potential was marked manually by a researcher, who scanned the entire EEG recording manually for all the participants in the active and sham groups, in the pre-stimulation and post-stimulation conditions. The number of eyeblinks in the length of recording was obtained and the eyeblink rate was calculated as the number of eyeblinks/minute. The same procedure was performed by a second researcher who was blinded to the conditions and the inter-rater validity was calculated. The average score was calculated from both independent researchers.

#### Saliva collection

Saliva was collected twice during each experiment: once immediately prior to NITESGON stimulation and once immediately after NITESGON stimulation as detailed in Experiment 1.

#### Procedure

Participants performed the auditory oddball task twice, once immediately before and once immediately after the NITESGON session. Saliva was also collected immediately before and immediately after the NITESGON session. Participants were randomly assigned to the active or sham NITESGON group. The researcher who controlled the NITESGON device was not involved in instructing the participant; this was performed by a second researcher who was blind to the stimulation protocol.

#### Statistics peak-to-peak P3b amplitude

EEG data was compared using a repeated measures ANOVA with groups (active vs. sham) and condition (deviant vs. standard) as the between-subjects variable, and the peak-to-peak amplitude before and after stimulation as the within-subjects variable. A simple contrast analysis was applied to compare specific effects using a Bonferroni correction.

#### Statistics sEBR

We conducted a repeated measures ANOVA with group (active vs. sham) as between-subjects variable, and the average eye blink rate before and after stimulation as within-subjects variable. A simple contrast analysis was applied to compare specific contrasts using a Bonferroni correction.

#### Statistics saliva

Using the saliva collected via the passive drool method, sAA levels were measured which were compared between the groups as detailed in experiment 1.

#### Statistics correlation

Pearson correlations were calculated between the difference in sAA, peak-to-peak P3b amplitude and sEBR before and after NITESGON stimulation.

### Experiment 8 – Dopamine

#### Participants

Participants were 20 right-handed adults (8 males, 12 females; mean age was 35.23 years, *Sd* = 2.63 years), half of who were selected due to their medical record indicating they were taking flupentixol (0.5mg)/melitracen (10mg)(Deanxit), a dopamine antagonist (i.e., D1 and D2)^89^, for their tinnitus at least two weeks prior to the onset of the study. The remaining participants were of matching age and gender with a similar educational background. Participants were screened and enrolled similar to Experiment 1.

#### Word-association task

Associative memory performance was measured using the same computerized Swahili-English verbal paired-associative learning task used in Experiment 1, however, the English words were replaced by Dutch words.

#### NITESGON

All participants received active NITESGON immediately following the word association task on visit 1 using the following parameters: a 5 second ramp-up period, followed by a constant current of 1.5 mA for 25 minutes, and finished with a 5 second ramp-down period.

#### Procedure

Eligible participants were scheduled for two visits to complete the study. Visit 1 consisted of the word-association task followed by active NITESGON stimulation. Participants were asked to refrain from studying or searching for the learned word pairs throughout the week. Participants returned 3 to 4 days after their first visit for memory testing to measure possible (long-term) effects on associative memory performance, but did not receive NITESGON, nor were they able to review word-pairs.

#### Statistics task

For visit 1 learning, a one-way ANOVA was conducted with the cumulative learning rate over the different study periods as the dependent variable and two groups (antagonist or no antagonist) as between-subjects variable. To look at the memory effect (recall) 7-days after learning, we applied a one-way ANOVA with group as the between-subjects variable and correctly recalled words as dependent variable.

#### Blinding

For Experiments 1-7, participants were asked to guess if they thought they were placed in the active or control group. A χ^2^ analysis was used to determine if there was a difference between what stimulation participants perceived in comparison what participants expected.

